# The critical role of *Dmnt1* during spermatogenesis is not predictable, but knockdown does cause pervasive differential transcription

**DOI:** 10.1101/2023.02.09.527865

**Authors:** Christopher B. Cunningham, Emily A. Shelby, Elizabeth C. McKinney, Robert J. Schmitz, Allen J. Moore, Patricia J. Moore

## Abstract

Cytosine methylation and its machinery influence the gene expression of many eukaryotes; however, insects are an exception to this general tenet despite many lineages retaining methyltransferases and methylated genomes. Here, we tested the *a priori* hypothesis that perturbed genetic pathways will be associated with meiosis using transcriptomics because previous our work using *Oncopeltus fasciatus* shows that gametogenesis is interrupted at meiosis following knockdown of *DNA methyltransferase 1* (*Dnmt1*). Testes, which are almost exclusively contain gametes at varying stages of development, were sampled at 7-days and 14-days following knockdown of *Dmnt1* using RNAi. Using microscopy, we found actively dividing spermatocysts were reduced at both sampling points. However, we found limited support of perturbation for our predicted cell cycle and meiotic pathways and only at 14-days. We found that Gene Ontology terms had no preferential enrichment for meiosis-associated genes. Following our *a priori* tests, we used the full dataset to uncover further candidate pathways influenced by *Dnmt1* knockdown. Very few genes were differentially expressed at 7-days, but nearly half were at 14-days. We did not find strong candidate pathways for how *Dnmt1* knockdown was achieving its effect through Gene Ontology term overrepresentation analysis. Given the evidence from microscopy, we propose *Dnmt1* knockdown results in condensed nuclei after mitosis-meiosis transition and then cellular arrest. This explanation posits that differential gene expression is a product of comparing healthy to arrested cells and is not a targeted response.

## Introduction

A complete evolutionary understanding of any phenotype requires understanding both its function – its action and how that influences fitness - and the mechanisms that underpin its variation – the substrate that evolution acts upon. As genetic technologies continue to expand, we can ask these questions of molecular phenotypes (Maleszka and Kucharski, 2022). Cytosine methylation, the addition of a methyl group to the nucleotide cytosine, is very important to cell function and survival (Law and Jacobsen, 2010; Schmitz et al. 2019). For many domains of life, cytosine methylation is an epigenetic mechanism that negatively regulates gene expression (Schmitz et al. 2019). In contrast, while the methylation levels within gene bodies across insect show a consistent association with highly and broadly expressed genes (Yan et al. 2014; Libbrecht et al. 2016; Duncan et al. 2022), differences of cytosine methylation are not associated with difference of gene expression of insects except for rare exceptions (Monardin and Brendel, 2021; Duncan et al. 2022). Cytosine methylation itself and its machinery are variably conserved across most insect lineages (Bewick et al. 2017; Provataris et al. 2018; Duncan et al. 2022) indicating that it has functional roles for insects. One consistent function of cytosine methylation machinery of insects is a role during gamete and embryo formation (Zwier et al. 2012; Schulz et al. 2018, Bewick et al. 2019; Amukamara et al. 2020; Gegner et al. 2020; Washington et al. 2021; Arsala et al. 2022). This role has been demonstrated through both pharmacological and genetic manipulations. For some species this role has been identified to be critical during meiosis specifically (*Oncopeltus fasciatus;* Amukamara et al. 2020, Washington et al. 2021). A mechanistic understanding of cytosine methylation and its machinery’s role is needed to address why it is critical during gamete formation (Bewick et al. 2019; Washington et al. 2021), despite not acting as a direct mechanism of gene regulation.

The large milkweed bug *Oncopeltus fasciatus* has become a model system for studies of evolution and development, especially of hemipteran insects (Chipman, 2017). There are also relatively high levels of cytosine methylation of *O. fasciatus*, as well as functional copies of the two DNA cytosine methyltransferase genes (Bewick et al. 2019). We have therefore been investigating the role of *Dnmt1* in this species. Specifically, we have investigated the function of the maintenance DNA methyltransferase responsible for replicating cytosine methylation patterns after a cellular division (Schmitz et al. 2019), *Dnmt1*, during gamete formation of both ovaries and testes. When adult female *O. fasciatus* are injected with *Dnmt1* RNA interference (RNAi), they fail to produce viable eggs and gamete production stops (Bewick et al. 2019). When injected during a juvenile stage prior to formation of primary oocytes in the developing ovary, knockdown of *Dnmt1* completely ablates gamete formation despite somatic ovaries developing through a normal trajectory (Amukamara et al. 2020). When *Dnmt1* is knocked down during a pre-meiosis stage of juvenile testis development, males emerge as adults with significantly smaller testes that have fewer developing sperm (Washington et al. 2021). Similar to females, when injected as adults, spermatogenesis is blocked and no further sperm develop resulting in reduced fertility once sperm that had been in the process of development at the time of treatment have been used up (Washington et al. 2021). These results indicate that the reduction of *Dnmt1* specifically affects production of gametes rather than gamete maturation. This led us to hypothesize that *Dnmt1* influences gametogenesis at meiosis.

With this study, we used RNA-seq to test the hypothesis that *Dnmt1* influences spermatogenesis by influencing the expression of meiosis genes. In females, this reduction of fecundity is associated with the differential gene expression of several hundred genes (Bewick et al. 2019). Similar to other insects, this differential gene expression is not associated with differential methylation (Bewick et al. 2019). However, if *Dnmt1* specifically affects the primary oocytes, which represent a small proportion of the cells within the adult ovary, it may be that any specific signatures of differential gene expression might be masked by the large number of unaffected somatic cells. The situation in the testis is very different. Each testis tubule is comprised of spermatocysts containing sperm at varying stages of development, and the majority of cells within the testis are spermatogonia, spermatocytes, or developing spermatids. Given that the underlying phenotype in males and females appears to be similar, a block to the transition from germ cell (the diploid oogonia or spermatogonia) to gametes, we compared the gene expression pattern of testes following *Dnmt1* knockdown using RNAi to control testes. To capture any early events as opposed to more global changes that might result from downstream impacts of blocked development not directly related to the knockdown of *Dnmt1*, we sampled testes at 7-days and 14-days post-injection, which are earlier than the 21-day post-injection samples taken in the Washington et al. (2021) where testes have significantly reduced numbers of developing spermatocytes and a highly disrupted structure. With these earlier sampling points, we aimed to capture the gene expression changes that were leading to the developmental arrest of spermatogenesis, presumably at the transition between diploid spermatogonia and haploid spermatocytes. We performed three analyses. First, we contrasted a set of *a priori* candidate genes at both sampling points between treatment and control samples. Second, we looked for the enrichment of specific GO terms at each sampling point; hypothesizing that we should see enrichment of meiosis, spermatogenesis, and gametogenesis and not of mitosis or cell cycle. Third, as an exploratory aim, we also compared the differential expression between the treatments at the gene and GO term level to identify new candidate genes and pathways influenced after *Dnmt1* knockdown. We found very little differential gene expression at 7-days even though we were able to detect phenotypic changes in the testes at this sampling point. In contrast, there was a very large number of differentially expressed genes at 14-days. None of the *a priori* candidate genes were differentially expressed at 7-days. Six of seventeen candidate genes were differentially expressed at 14-days, but did not concentrate on any one pathway. With our *a priori* GO term enrichment analysis, we found all terms were depleted at 7-days (i.e., genes with those GO terms were preferentially among the least differentially expressed genes). At 14-days, no GO terms were enriched or depleted. For our exploratory analysis to find new candidate pathways, at 14-days, many pathways were enriched among the differentially expressed genes, but they did not center around one general cellular process or phenotype. These results suggest a rapid and highly perturbed transcriptional environment in the absence of *Dnmt1* that leads to cellular arrest among spermatocysts as they attempt to progress through gametogenesis at the meiosis stage. Previous experiments provide no evidence that *Dnmt1* is acting on the regulation of gene expression itself. We therefore propose an alternative model of how the maintenance *Dnmt1* perturbation might affect cell function and lead to a block to the development of spermatogonia into spermatocytes.

## Methods and Material

### Overview

To understand the functional role that *Dnmt1* plays during gamete formation of male *O. faciatus*, we used RNAi to knockdown *Dnmt1* gene expression at two sampling points post-treatment: 7-days and 14-days. After the treatment at two sampling points, we used confocal microscopy to phenotype the testes and determine the state of spermatocytes. We then profiled gene expression using RNA-seq of testes at the same two sampling points to understand how *Dnmt1* knockdown perturbed the testes transcriptome.

### Animal Colony and Husbandry

*Oncopeltus fasciatus* colony stocks were purchased from Carolina Biologicals (Burlington, NC). The colonies were maintained in Percival incubators under a 12:12 hour light:dark cycle at 27°C. The animals were fed organic raw sunflower seeds and had *ad libitum* deionized water. We needed to collect animals of known age so we removed eggs from the colonies and housed them in plastic storage containers with food and water. Nymphs were sexed at the fourth instar and separated into single sex colonies. Containers were checked daily for adult eclosions. New adults were placed into individual petri dishes with food and water.

### RNA Interference (RNAi) Synthesis, Administration, and Quality Control

We created RNAi constructs of *Dnmt1* following Bewick et al. (2019). Briefly, DNA templates of *Dnmt1* were prepared via PCR. Following that, double stranded RNA was synthesized and then digested with an Ambion MEGAscript kit (ThermoFisher Sci, Waltham, MA) following the manufacturer’s protocol. This reaction was purified with a phenol:chloroform:IAA extraction followed by a sodium acetate precipitation. Concentration was measure by a Qubit using the ssRNA kit following the manufacturer’s protocol. Sense and anti-sense strands were then allowed to anneal. We used *eGFP* as an exogenous control construct. Before administration, the RNAi construct concentration was adjusted to 4 μg/μL in injection buffer (5 mM KCl, 0.1 mM NaH2PO4) for both *Dnmt1* and *eGFP*. Sexually mature, virgin males (~7 days post-adult ecolsion) were injected with 3 μL ds-*Dnmt1* RNA or *eGFP* using a pulled capillary needle between the third and fourth abdominal segments (Chesebro et al. 2009). Males were paired with a virgin female to allow for mating and stimulate sperm production. Pairs were kept in petri dishes under standard rearing conditions until they sampled. Males were haphazardly allocated to treatment group.

We have established previously established there is no difference between controls using RNA constructs with no specificity to *O. fasciatus* genome sequence (e.g., *eGFP*) and buffer alone injections (Bewick et al. 2019). Therefore, we did not include a buffer alone control. Previously, we have targeted RNAi constructs to two different regions of *Dnmt1*; the cytosine-specific DNA methyltransferase replication foci domain (RFD) and the DNA methylase domain (AdoMet). These give rise to identical phenotypes, which suggests minimal off-target effects (Bewick et al. 2019). Thus, here, we only targeted against the RFD consensus domain.

### RNAi validation by Quantitative RT-PCR (qRT-PCR)

We assessed the effectiveness of *Dnmt1* knockdown, using qRT-PCR. We synthesized cDNA using 500 ng RNA with qScript cDNA SuperMix (Quanta Biosciences, Gaithersburg, MD). Validated primers were used from Bewick et al (2019). GAPDH was used as the endogenous reference gene. We used Roche LightCycler 480 SYBR Green Master Mix with a Roche LightCycler 480 (Roche Applied Science, Indianapolis, IN) with 3 technical replicates of 10 μL reactions as previously reported (Cunningham et al. 2014). We used the ΔΔ*C*_T_ method to estimate differences of expression using *eGFP* samples as our comparison group (Livak and Schmittgen, 2001).

### Phenotypic analysis of *Dnmt1* knockdown

Virgin, adult males were collected on the day of emergence. Males were injected with *ds-eGFP* or ds-*Dnmt1* as described above at 7 – 10 days post-adult emergence. Males were haphazardly assigned to two dissection days. One group of males were dissected 7 days post RNAi treatment and a second group was dissected 14 days post RNAi treatment.

Testes were dissected from the males in PBS and processed for microscopy as described above. To assess the activity of early-stage spermatocytes, the testis tubules were stained with anti-phosphohistone H3 Ser 10 antibody (pHH3; Millipore antibody 06-570, Sigma Aldrich, St. Louis, MO). Anti-pHH3 stains for chromosome condensation in preparation for mitosis and meiosis (Hans and Dimitrov 2001, Prigent and Dimitrov, 2003). Primary antibody staining was visualized with a secondary antibody, Alexa Fluor goat-anti-rabbit 647 (ThermoFisher Scientific, Waltham, MA). Testis tubules were counterstained with DAPI (0.5 μg/mL PBT) to visualize nuclei. Testis tubules were visualized on a Zeiss 710 confocal microscope or an EVOS (ThermoFisher Scientific, Waltham, MA).

To quantify the number of actively dividing spermatocysts, 20 testis tubules from a minimum of ten males per treatment were examined. Anti-pHH3 antibody stains both spermatogonia undergoing mitotic divisions and spermatocysts undergoing meiotic divisions (Figure 1A). The numbers of positively stained spermatocysts in either control (ds-*eGFP*) or knockdown (ds-*Dnmt1*) males were compared at within each sample using a *t*-test with Welch’s correction for unequal variance and the prediction that ds-*Dnmt1* samples would have fewer dividing cells.

**Figure 1.**
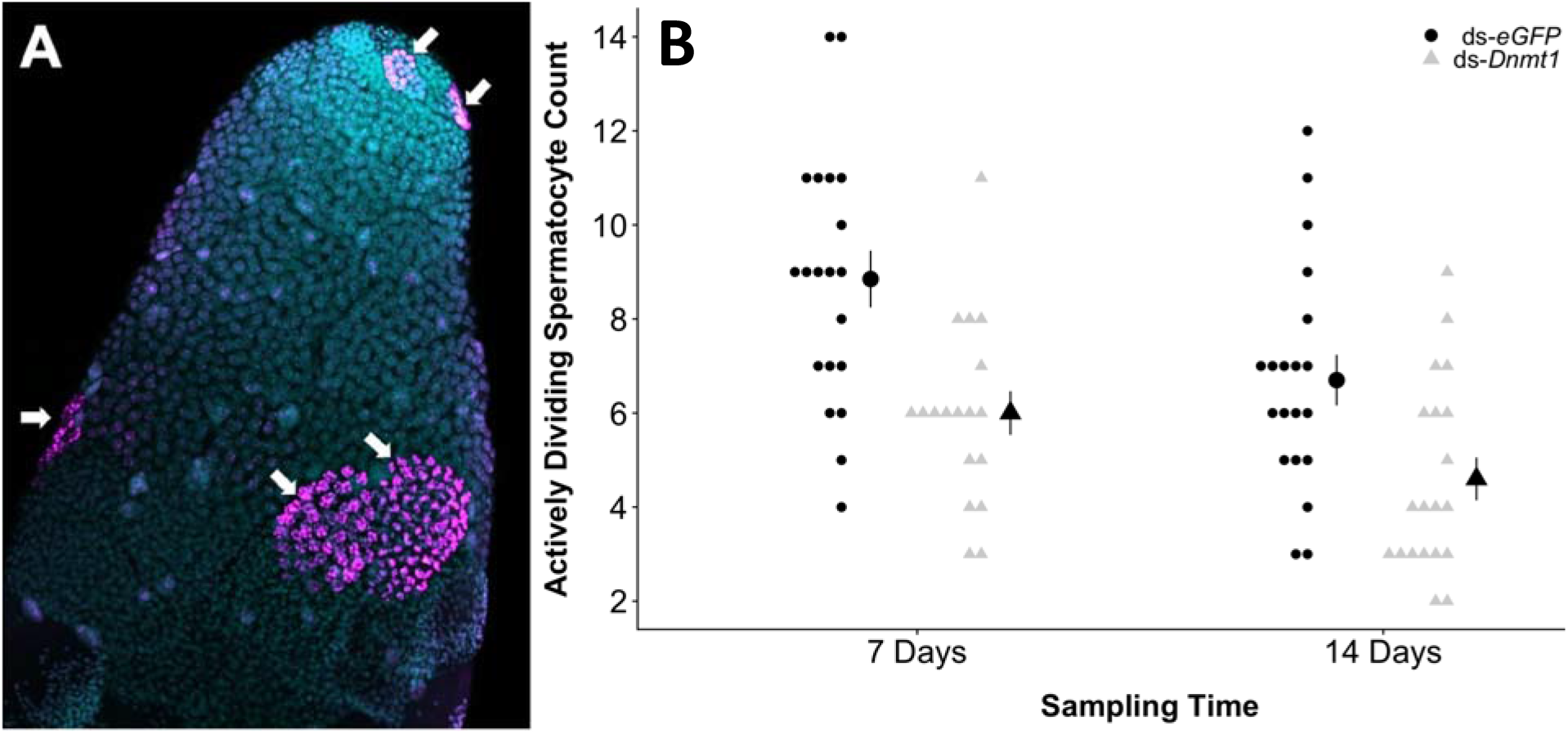
Male gamete formation is highly perturbed following knockdown of *Dnmt1* revealed by testes tubules stained with anti-phosphohistone H3 antibody. (A) Representative image of a testis tubule stained with anti-pHH3 antibody and DAPI. The spermatocysts that were in the act of dividing at the time of dissection are labeled with magenta (arrows). 20X magnification. (B) There were fewer dividing cells in *Dnmt1* knockdown males at both sampling times post-RNAi treatment compared to controls.

### RNA Extraction

The testes of males of each treatment were dissected out in ice-cold 1X PBS at the appropriate sampling point, flash frozen with liquid nitrogen, and stored at –□80°C until nucleotide extraction. Total RNA was extracted using a Qiagen Allprep DNA/RNA Mini Kit (Qiagen, Venlo, The Netherlands) following the manufacturer’s protocol. Homogenization of the testes were performed with a handheld Kimble pellet pestle in RLT buffer. Quantification was done with a Qubit fluorometer using the RNA BR kit.

### RNA-seq High-throughput Library Preparation & Sequencing

The extracted RNA of *O. fasciatus ds-Dnmt1* and control (ds-*eGFP*) biological replicates at 7- and 14-days post-injection was used to construct poly-A selected Illumina TruSeq Stranded RNA LT Kit (Illumina, San Diego, CA) following the manufacturer’s instructions with limited modifications. The starting quantity of total RNA was adjusted to 1.3 μg, and all reagent volumes were reduced to a third of the described quantity. We targeted 10M 2 × 150 bp read pairs per biological replicate using an Illumina NextSeq 500 with v3.1 chemistry. Removing libraries that were not viable left 13 7-day *eGFP*, 10 7-day *dnmt1*, 15 14-day *eGFP*, and 14 14-day *dnmt1*. Libraries were sequenced at the Georgia Genomics and Bioinformatics Core (Athens, GA, USA).

### Read Quality Control and Mapping

Reads were initially assessed for quality with fastQC (v0.11.9; default settings; Andrews, 2010) to establish a baseline. Reads had adapters trimmed with cutadapt (v2.8; --trim-n -O 3 -u -2 -U -2 -q 10,10 -m 30; Martin, 2011) using the TruSeq adapter sequences. Reads were again assessed with fastQC with default settings. Overlapping reads were combined with FLASh (v 1.2.11; default settings; Magoc and Salzberg, 2011). As a final QC step, reads that mapped to rRNA genes were removed with SortMeRNA (v4.3.3; all gff entries annotated as rRNA; Kopylova et al. 2012).

We used the NAL i5k *O. faciatus* genome (the “BCM-After-Atlas” version) & annotation (Official Gene Set v1.2; Panfilio et al. 2019). These were downloaded from the NAL i5k site: https://i5k.nal.usda.gov/content/data-downloads. This was the most current version of the genome and gene annotation at the time of analysis. HISAT2 (v2.2.1; no soft-clipping; Kim et al. 2019) was used to map reads to the genome (Supplementary File 1). Extended reads (i.e., reads that were combined by FLASh) had ~30% higher mapping rate. Mappings were converted to read counts by StringTie (v2.1.7; Pertea et al. 2015) following the manual instructions for export to DESeq2.

### Functional Annotation

We updated the functional annotation of the *O. faciatus* proteome/transcriptome using eggNOG-mapper webserver (http://eggnog-mapper.embl.de/; v 2.19; Cantalapiedra et al. 2021). This annotated 12,526 of 19,615 gene models (63.8%) with Gene Ontology terms, which allowed us to perform a GO term enrichment analysis and a GO term overrepresentation analysis.

### Differential Gene Expression

Read counts were imported into R using tximport (Bioconductor v1.20.0; Soneson et al. 2015) following the manual’s instructions. We used R (v4.1.0; R Core Team, 2021) within an RStudio IDE (build 492; RStudio Team, 2022) for the differential gene expression analysis.

DESeq2 (Bioconductor v1.32.0; default settings; Love et al. 2014) was used for all DGE analyses following the programmers’ suggestions for exploratory data analysis and sample/analysis quality control. *eGFP* was set and used as the comparison group for all analyses. 7-day and 14-day samples were analyzed separately as previously discussed to generate a list of differentially expressed genes between control and experimental treatments. The model matrix specification was also checked to ensure correct specification (i.e., that program was contrasting the samples in the correct way). After importing and initial analysis, samples were plotted with a PCA to visually check for outliers according to the manual’s recommendation. One 7-day *Dnmt1* sample (1.30) was removed. Two 14-day samples were also removed – one *Dnmt1* (1.32) and one *eGFP* (1.9) remaining.

After removal of the outliers each analysis was repeated using the same settings of DESeq2 and the results were again quality controlled. We used the default dispersion estimator and shrinkage method, apeglm (Zhu et al. 2019). We used s-values to estimate statistical significance after false discovery rate correction at the level of 0.05 (Stephens 2016). Results were visualized with the fviz_pca_ind function of the factoextra R package (Kassambara and Mundt 2020).

We also compared the overlap of the differentially expressed genes here to the differentially expressed genes of *O. fasciatus* ovaries under the same treatment scheme using the intersect function of R. A simulation analysis was conducted to see if any overlap observed was statistically significantly enriched or depleted (Cunningham et al. 2019).

### *a priori* Candidate Gene Screen

The first test of our hypothesis was to assess the influence *Dnmt1* knockdown had on a series of candidate genes. These genes were chosen based on literature searches for genes involved in meiosis and spermatogenesis of insects. These include cell fate and proliferation genes that we did not expect to be differentially expressed (members of the *Wnt* and *Frizzled* families), cell cycle control genes some of which we did expect to be differentially expressed (*Vasa*) and some of which did not expect to be differentially expressed (*Cyclin B3, Cyclin D2, Cyclin D3, Cell Division Cycle 20, Cell Division Cycle 25* family members), maintenance of chromosome genes that we did expect to be differentially expressed (*Structural Maintenance of Chromosome 3* family members), meiotic transition gene that we did expect to be differentially expressed (*Boule*), and meiotic recombination genes that we did expect to be differentially expressed (*SPO11 Initiator of Meiotic Double Stranded Breaks, MutS Homolog 5, Meiotic Nuclear Divisions 1, Homologous-Pairing Protein 2*). None of the candidates had specific directional changes, except decrease expression of *Vasa* and *Boule* within the Dnmt1 knockdowns. We extracted the raw P values for expression differences from the DESeq2 results and compared them with FDR corrected P values we generated using the Benjamini-Hockberg procedure (Benjamini and Hockbeg 1995)

### *a priori* GO Term Enrichment Analysis

As secondary test of our hypothesis, we selected five high-level GO terms that represented the cellular processes that we thought were being perturbed or not after *Dnmt1* knockdown. These were GO:0007049 Cell Cycle, GO:0000278 Mitotic Cell Cycle, GO:0051321 Meiotic Cell Cycle, GO:0007276 Gametogenesis, and GO:0007283 Spermatogenesis. With our previous observed phenotypes and the one observed with the microscopy here, we expect Meiotic Cell Cycle, Gametogenesis, and Spermatogenesis to be enriched. We expected Cell Cycle and Mitotic Cell Cycle not to show enrichment. We included GO:0070265 Necrotic Cell Death and GO:0006915 Apoptotic Process to assess if cells were undergoing these processes or if they had simply arrested. We used S values from DESeq2 to rank order the expressed genes at 7- and 14-days separately. We used fgsea R package (Korotkevich et al. 2021) with default parameters, except allowing 1,500 genes as a maximum. Results were visualized with the same R package.

### Gene Ontology (GO) Term Overrepresentation Analysis

We found overrepresented terms to give us biological processes that might be perturbed using the topGO package of R (v2.48.0; Alexa et al. 2006). This analysis is often termed an enrichment analysis (Huang et al. 2009), but statistically it tests if there is an overrepresentation of a GO term from a list of interesting genes (usually those that are differentially expressed) compared with the expectation of the same number of random picks. It does not calculate if a specific GO term (or another type of annotation; e.g., KEGG) is enriched at the beginning of an ordered gene set. We performed GO term tests using Fisher’s exact test with the weighted algorithm. GO terms of genes of interest were compared to all expressed genes within our samples. Updated GO terms can be found in Supplementary File 2.

To refine our overrepresented GO terms to even higher-level summaries of the processes involved and identify central processes enriched from our treatments, we performed a semantic similarity analysis using REVIGO using default settings (Supek et al. 2011).

## Results

### *Dnmt1* RNAi Knockdown

*Dnmt1* was effectively knocked down in experimental males treated with ds-*Dnmt1* RNA. Using qRT-PCR, we confirmed *Dnmt1* was downregulated in the ds-*Dnmt1* samples (7 day: mean fold expression difference = −0.677, *t*_24_ = 7.250, P = 8.609e-8; 14 day: mean fold expression difference = −0.679, *t*_25_ = 4.288, P = 0.00012). For both datasets (7-day, 14-day) the *5’* end of *Dnmt1* (OFAS015351) was statistically significantly downregulated in *Dnmt1* RNAi treatment (Table 1). For neither dataset was the 3’ end of *Dnmt1* (OFAS018396) statistically significantly differentially expressed, but both were downregulated in the *Dnmt1* RNAi treatment (Table 1).

**Table 1.** *a priori* candidate genes tested for differences between 7- and 14-day Control and *Dnmt1* knockdown treatments. Values were standardized using the vst function of DESeq2. Bolded values are statistically significant after Benjamini-Hockberg FDR Correction.

| Gene Name | Accession | Pathway | 7-days |  |  | 14-days |  |  |
| --- | --- | --- | --- | --- | --- | --- | --- | --- |
|  |  |  | Standardized Expression | Fold Change | FDR-corrected P value | Standardized Expression | Fold Change | FDR-corrected P value |
| Dnmt1_5' | OFAS015351 | Cytosine methylation | **8.5** | **-0.369** | **3.12E-04** | **6.8** | **-1.355** | **3.90E-12** |
| Dnmt1_3' | OFAS018396 | Cytosine methylation | 7.2 | -1.47E-06 | 0.829 | 5.7 | -0.389 | 0.205 |
| Wnt-like | OFAS019215 | Cell fate & proliferation | NA | NA | NA | NA | NA | NA |
| Wnt-like | OFAS025063 | Cell fate & proliferation | 7.1 | 1.25E-06 | 0.994 | 5.5 | 0.142 | 0.621 |
| Frizzled-like | OFAS012753 | Cell fate & proliferation | 7.3 | -4.82E-06 | 0.037 | 5.7 | -0.302 | 0.226 |
| Frizzled-like | OFAS013300 | Cell fate & proliferation | 9.5 | -7.16E-06 | 0.245 | 7.8 | 0.047 | 0.766 |
| Frizzled-like | OFAS013301 | Cell fate & proliferation | 7.8 | 1.03E-06 | 0.955 | 6.3 | -0.036 | 0.875 |
| Vasa | OFAS025106 | Germ cell Development | 10.7 | -3.68E-06 | 0.663 | **8.6** | **-0.622** | **0.013** |
| CYCB3 | OFAS004926 | Cell cycle control | 7.9 | 1.29E-06 | 0.618 | 5.8 | -0.337 | 0.268 |
| CDC20 | OFAS011085 | Cell cycle control | 12.9 | 1.07E-05 | 0.853 | **10.7** | **-0.743** | **7.47E-05** |
| CDC25-like | OFAS004022 | Cell cycle control | 9.0 | -4.22E-05 | 0.336 | **7.0** | **-0.575** | **0.004** |
| CDC25-like | OFAS004025 | Cell cycle control | 7.1 | -5.29E-05 | 0.938 | 5.5 | 0.027 | 0.925 |
| CYCD2 | OFAS016512 | Cell cycle control | NA | NA | NA | NA | NA | NA |
| CYCD3 | OFAS016513 | Cell cycle control | 6.4 | 2.72E-06 | 0.149 | 5.2 | 0.384 | 0.178 |
| SMC3-like | OFAS014329 | Chromosomal Structure Maintenance | 7.9 | 1.43E-05 | 0.111 | 5.9 | -0.204 | 0.475 |
| SMC3-like | OFAS014330 | Chromosomal Structure Maintenance | 6.7 | -7.77E-05 | 0.971 | 5.1 | -0.549 | 0.092 |
| Boule | OFAS007982 | meiotic G2/M transition | 12.3 | -1.98E-06 | 0.121 | **10.2** | **-0.573** | **0.013** |
| SPO11 | OFAS017698 | Meiotic Recombination | 9.2 | 8.39E-07 | 0.777 | 7.1 | -0.453 | 0.052 |
| MSH5 | OFAS011763 | Meiotic Recombination | 6.5 | 1.18E-05 | 0.319 | **5.2** | **1.916** | **0.001** |
| MND1 | OFAS003294 | Meiotic Recombination | 9.6 | -9.14E-05 | 0.220 | 7.5 | -0.258 | 0.157 |
| HOP2 | OFAS000825 | Meiotic Recombination | 8.3 | 2.24E-05 | 0.032 | 6.7 | 0.346 | 0.088 |

### Microscopy – Effect of *Dnmt1* RNAi Knockdown

Knockdown of *Dnmt1* had a measurable phenotype in males sampled at both 7- and 14-days post injection. Knockdown of *Dnmt1* reduced the number of actively dividing spermatocysts (Figure 1A). The number of spermatocysts that were positively stained with anti-pHH3 antibody, which indicates cell undergoing mitotic or meiotic division, was reduced in males at both 7 days post-knockdown (mean difference = −3.45, *t*_37_._768_ = 4.098, P = 0.0001; Fig 1B) and 14 days post-knockdown (mean difference = −2.1, *t*_36,979_ = 2.978, P = 0.0026; Figure 1B).

### *a priori* Candidate Genes

There were no differentially expressed *a priori* candidate genes at 7-days (Table 1). Six of seventeen were at 14-days (Table 1). However, they did not concentrate around a particular process. Three of six cell cycle control genes were differentially expressed, while two of five meiotic genes were differentially expressed.

### *a priori* GO Term Enrichment

None of the GO terms were enriched at 7- or 14-days after *Dnmt1* knockdown. In fact, contrary to our expectation, all of the five GO terms that we made predictions for were statistically significantly absent from the top of the list of genes rank-ordered by difference of expression at 7-days (i.e., they were statistically significantly preferentially found at the bottom of the list of genes rank-ordered by their expression differences), while none were overrepresented at either the top or bottom of the rank-ordered list at 14-days (Fig. 2). The necrotic and apoptotic GO terms were not enriched at either sampling point.

**Figure 2.**
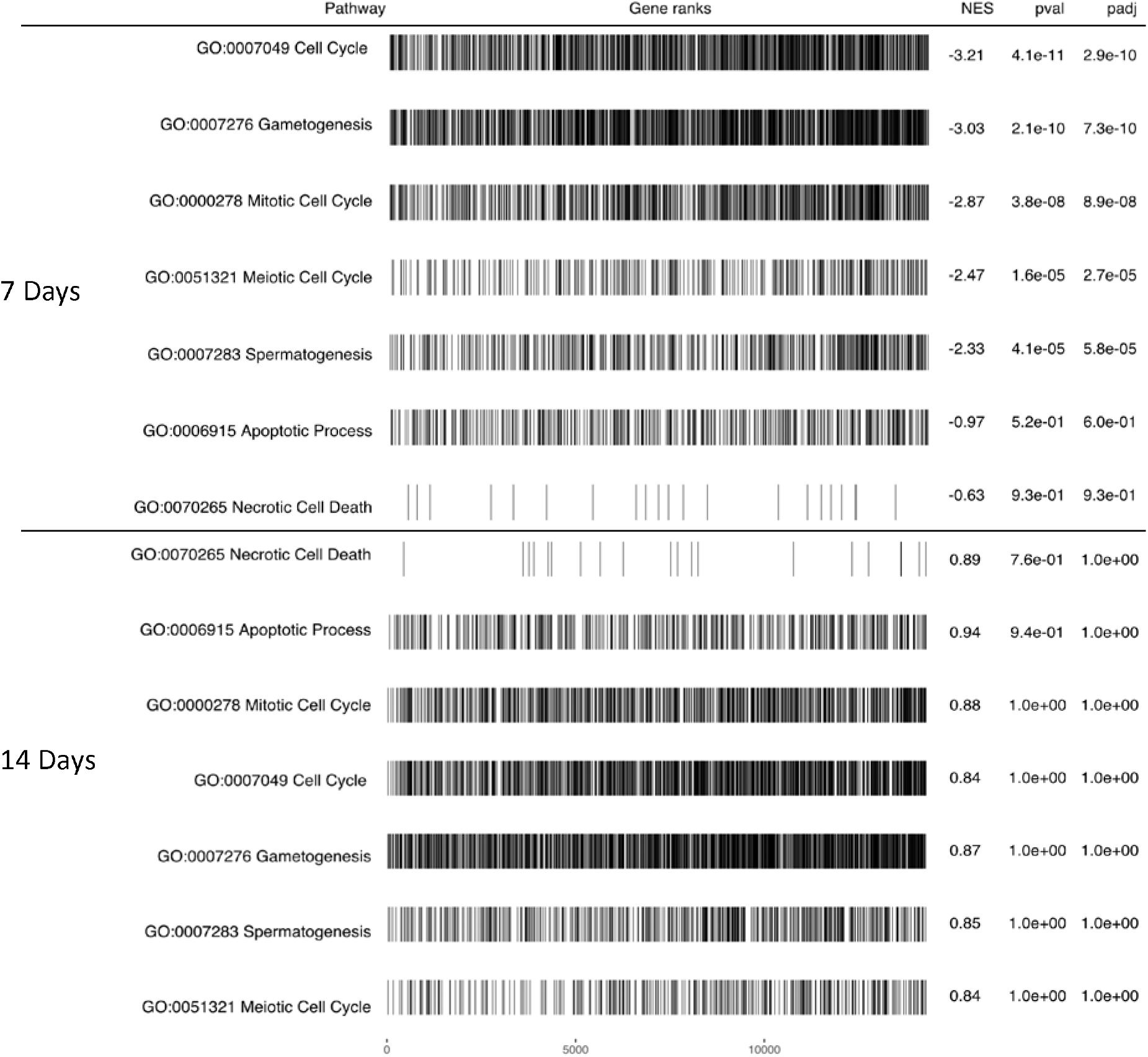
*a priori* GO terms move from depleted at 7-days to neither enriched nor depleted at 14-days. The genes collectively moved up the ordered differentially expressed gene, but not at a preferential rate. This aligns with a general, global response to the loss of *Dnmt1* not heavily, preferentially targeting one or a few pathways. NES = normalized enrichment score, pval = P value, padj = FDR-corrected P value.

### Differential Gene Expression

To identify the genes that are differentially regulated by the knockdown of *Dnmt1* that leads to the systematic depletion of spermatocysts among adult *O. fasciatus* testes, we investigated gene expression at two sampling points −7- and 14-days – post-injection. The individual treatments clustered together well at both sampling points (Fig. 3). The treatments are not differentiated at 7 days (Fig. 3A; Supplementary File 3), are but highly differentiated at 14 days (Fig. 3B; Supplementary File 4). There were 74 genes that were differentially expressed between the control treatment and the ds-*Dnmt1* treated samples at 7-days post-injection (23 up-regulated; 51 down-regulated). There were 6,746 genes that were differentially expressed between the control treatment and the ds-*Dnmt1* treated samples at 14 days post-injection (2,761 up-regulated; 3,985 down-regulated).

**Figure 3.**
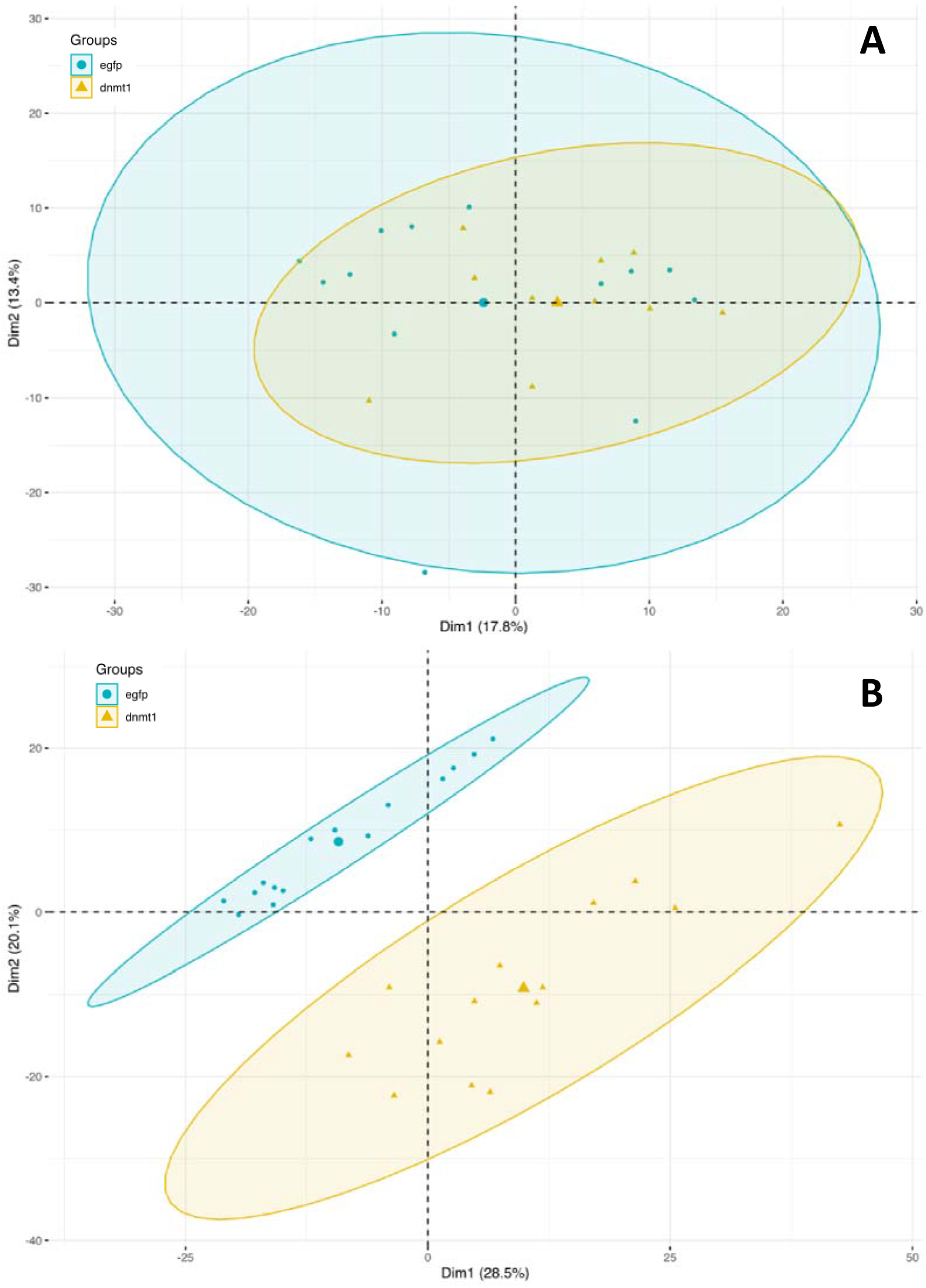
Gene expression knockdown of *Dnmt1* leads to increasing differential gene expression as testes initiate and progress through gamete formation after treatment. Here, samples are visualized with a Principal Component Analysis visualization. (A) Samples 7 days post treatment are highly overlapping and not differentiated by many gene expression differences. (B) Samples 14 days post treatment are highly non-overlapping and differentiated by many gene expression differences (~45% of expressed genes are different between the treatments at this sampling point).

The overlap between the differentially expressed genes found here and those of Bewick et al. (2019) using the same treatment against ovaries, was four for the 7-day list and 96 for the 14-day list. Both of these overlaps are statistically significant. The 7-day comparison had more than expected (mean simulated overlap = 1.32 P = 0.041) and the 14-day comparison had fewer than expected (mean simulated overlap = 119.24, P = 0.0022).

### Gene Ontology (GO) Term Overrepresentation

Among the differentially expressed genes between *Dnmt1* knockdown and *eGFP* control at 7-day samples, 113 GO terms were statistically significantly overrepresented. The top three Biological Processes terms with more than one gene were purine nucleobase biosynthetic process – GO:0009113, skeletal muscle organ development – GO:0060538, and mitotic sister chromatid cohesion – GO: GO:0007064 (P = 0.0014, 0.0016, 0.006, respectively); Molecular Function terms were calcium-dependent phospholipid binding – GO:0005544, cation binding – GO:0043169, and ribose phosphate diphosphokinase activity (P = 0.0017, 0.008, 0.0083, respectively); and for Cellular Compartments were nuclear meiotic cohesin complex - GO:0034991, Parkin-FBXW7-Cul1 ubiquitin ligase complex - GO:1990452, and integral component of Golgi membrane - GO:0030173 (P = 0.0087, 0.0087, 0.0125, respectively). Very few GO terms collapsed under a parent GO term in our semantic similarity analysis trying to understand what broad process was most perturbed with *Dnmt1* knockdown. A complete term list can be found in Supplementary File 5.

For the 14-day samples, 322 GO terms were statistically significantly overrepresented. The top three Biological Processes terms with more than one gene were response to unfolded protein – GO:006986, double-strand break repair via nonhomologous end joining – GO:006303, and mRNA splice site selection – GO:0006376 (P = 0.00011, 0.00016, 0.00019, respectively), Molecular Function terms were nucleoside-triphosphatase activity – GO:0017111, single-stranded DNA binding – GO:0003697, and electron transfer activity – GO:0009055 (P = 0.00014, 0.00038, 0.00062, respectively), and for Cellular Compartments were ribonucleoprotein granule – GO:0035770, CCR4-NOT core complex – GO:0030015, sperm flagellum – GO:0036126 (P = 0.00015, 0.00025, 0.00025, respectively). Very few GO terms collapsed under a parent GO term in our semantic similarity analysis trying to understand what broad process was most perturbed with *Dnmt1* knockdown. A complete term list can be found in Supplementary File 6.

For the overlap between the differentially expressed genes found here and those of Bewick et al (2019) for the 14-day samples, we found 168 GO terms. The top three Biological Processes terms with more than one gene were protein K63-linked deubiquitination - GO:0070536, melanocyte differentiation - GO:0030318, and negative phototaxis (P = 0.0002, 0.00025, 0.00032, respectively), Molecular Function GO terms were proteasome binding - GO:0070628, alditol:NADP+ 1-oxidoreductase activity - GO:0004032, and estradiol 17-beta-dehydrogenase activity - GO:0004303 (P = 0.00012, 0.00029, 0.000221, respectively), and for Cellular Compartments were molybdopterin synthase complex - GO:0042629, apical cortex - GO:0045179, and outer dynein arm – GO:0036157 (P = 0.00017, 0.00691, 0.00072, respectively). Very few GO terms collapsed under a parent GO term in our semantic similarity analysis. A complete term list can be found in Supplementary File 7.

## Discussion

Our previous work on *O. fasciatus* shows that the machinery for cytosine methylation is needed for successful gamete production (Bewick et al. 2019; Amukamara et al. 2020; Washington et al. 2021). However, the genetic underpinnings of this effect remain poorly explained. Here, we reduced the gene expression of *Dnmt1* in sexually mature *O. fasciatus* males. Following treatment with RNAi, we sampled experimental male testes at two sampling points, 7- and 14-days post treatment, which is before a large reduction of spermatocytes and testes size is observed. Spermatocysts at both sampling points have reduced cell division at intermediate levels to what is seen to a later sampling point of 21 days (Washington et al. 2021). Despite the observation that downregulation of *Dnmt1* already affected spermatogonial division rates, none of the *a priori* candidate genes were differentially expressed at 7-days, although we did see differential expression for a subset at 14-days. None of these differentially expressed genes concentrated within one molecular pathway. The *a priori* GO term enrichment showed depletion of all terms at 7-days and no enrichment of any at 14-days. With our secondary analysis to discovery more potential candidate pathways influenced by *Dnmt1* knockdown, we found few gene expression difference at 7-days, but a large amount at 14-days post treatment (~45% of expressed genes are differentially expressed). There was no clear consensus among the overrepresented GO terms to suggest how this cellular arrest is being achieved. Instead, we propose that the differential gene expression seen is being driven by a healthy vs arrested cell contrast and that the effect of a reduction of *Dnmt1* expression is a general, global response tied to chromatin condensation, not a set of specific genomic loci that are being targeted. It remains an open question why following a reduction *Dnmt1* expression (and therefore cytosine methylation after a cellular division) cell arrest with highly condensed nuclei.

Our hypothesis did receive strong support from either a screen of candidate genes or GO term enrichment of predicted pathways. Given the phenotype observed here and previous studies, we predicted meiosis, and its molecular pathways would be highly perturbed, giving rise to a loss of meiotic progression and a cessation of spermatocyte production observed. The *a priori* candidate gene screen did not reveal any molecular pathway being overwhelmingly affected. Both the collection of cell cycle and meiotic genes contained differentially expressed genes, but it was not the majority of either group; 3/6 for cell cycle and 2/5 for meiotic genes. These results align with the observation that cells appear to arrest at the meiotic stage and do not progress through further cell division which would require completely cell cycles.

Additionally, as with the candidate gene analysis, our hypothesis received little support from our GO term enrichment analysis. None of the seven GO terms that we tested were enriched at 7-days (i.e., the genes with those GO terms were not preferentially found at the beginning of the rank-ordered gene list). In fact, the GO term searched were preferentially found at the end of the gene list at the 7-day mark, except for necrotic and apoptotic processes which did not have statistically significantly pattern (i.e., were among genes that showed the smallest differences of expression between the control and *Dnmt1* knockdown samples). Like our candidate gene results, this suggests that few pathways were perturbed at this sampling point and the is no evidence that the cells are aborting themselves. At 14-days, none of the GO terms were enriched nor were they preferentially found at the end of the rank-ordered gene list. This means that these genes collectively increased their rank within the rank-ordered gene lists based on expression differences from 7-days to 14-days, but not at a rate that made them statistically enriched at the top of the 14-day gene list. The necrotic and apoptotic GO terms were not enriched at either time point suggesting that cells were not actively promoting either of these processes at either sampling point. Taken together, we suggest this supports the hypothesis that the gene expression difference at 14-days is a global response to *Dnmt1* knockdown and is not highly targeted to any specific set of molecular pathways. Additionally, the cells do not have a detectable signal that they are undergoing any process to actively remove themselves from the cell population.

At the broadest level, our RNA-seq results suggest that testes with reduced *Dnmt1* expression have an increasingly degradation of their transcriptional environment. The 74 differentially expressed genes of the 7-day samples did not contain many transcription factors or developmental pathway genes as we expected. At 14-day nearly half of all the expressed genes are differentially expressed – 6,794 of 14,984 genes (45.3%). Within ovaries treated in the same way and sampled at 10-days post treatment, there were an intermediate amount of 236 differentially expressed genes (Bewick et al. 2019). There was little overlap from our 7-day differentially expressed gene list with the ovary samples and fewer than statistically expected with the 14-day list.

Our differentially expressed genes lists did not produce strongly overrepresented GO terms where many were associated around a consensus biological pathway or process; all of the top five GO terms produced by the semantic similarity reduction for any three of the categories for both days were single standalone terms. While this might be expected for the 7-day samples with few differentially expressed genes, the 14-day samples have more than enough information to uncover any strongly disrupted biological process. The overlap between Bewick et al. (2019) and our samples did not produce any strong candidate pathways for *Dnmt1*’s influence after reduction either. These pieces of evidence led us to suggest that gametes are proceeding along a developmental pathway until *Dnmt1* is needed, and when *Dnmt1* is not present due to gene knockdown, the cell has a massive, pervasive, and non-specific disruption of its transcription environment.

We propose that *Dnmt1*’s action during gametogenesis influences chromosomal dynamics and nuclei condensation. We propose this idea based on multiple and repeatedly seen lines of evidence. This mechanism of action would explain why a reduction of *Dnmt1* expression 1) leads to a highly perturbed transcriptional environment but why cytosine methylation is not associated with gene expression directly, 2) why *Dnmt1* RNAi tissues have high condensed nuclei at the boundary where spermatocysts usually enter the first meiotic division, and 3) why *Dnmt1* has a specific and critical role during meiosis. Gene expression knockdown of *Dnmt1* does lead to reduced cytosine methylation (Bewick et al. 2019; Amukamara et al. 2020; Washington et al. 2021). However, cytosine methylation differences are not directly causal for differential gene expression for *O. fasciatus* (Bewick et al. 2019), or for insects generally (Duncan et al. 2022). When gene expression of *Dnmt1* is reduced hundreds to thousands of gene are differentially expressed for both male and female reproductive tissues (Bewick et al. 2019; this study). We posit that the differential gene expression seen between *Dnmt1* gene expression knockdown and control samples is attributable to comparing samples where one set is naturally progressing through gamete formation while the other is in a highly arrested state. We do not believe that this arrest is an active process the cells are undergoing, but rather is a function of the cell not being able to progress. This aligns with there being no signal from the GO term enrichment the cells are undergoing any cellular death process; necrotic or apoptotic; and that the cell looks healthy at a gross anatomical level. Even though differences of methylation of individual genomic loci do not lead to differences of gene expression between treatments, high cytosine methylation is associated with high gene expression between different loci (Yan et al. 2014; Libbrecht et al. 2016; Gladstad et al. 2016). This trend is particular to insects in contrast to plants, fungi, and vertebrates for which high cytosine methylation is often associated with heterochromatin (Schmitz et al. 2019). Highly condensed nuclei are seen for testis tubules with gene expression knockdown of *Dnmt1* (Washington et al. 2021). This effect is most pronounced at the region where spermatocysts transition from spermatogonia to spermatocytes through meiosis (Washington et al. 2021). When *Dnmt1* gene expression is reduced by RNAi, it also leads to a reduction of cytosine methylation of somatic tissues (Amukamara et al. 2020). However, this reduction does not impact life span, suggesting that the lack of cytosine methylation within somatic tissues does not have a strong impact on the somatic cells of the organism (Amukamara et al. 2020). This also points to a difference between somatic and reproductive cells which aligns with the presence and absence of meiosis. What remains to be directly explained by this model is why reduced *Dnmt1* gene expression knockdown or a lack of cytosine methylation is associated with highly condensed chromatin (or conversely an inability to de-condense chromatin) as gametogenesis progresses from earlier to later stages.

This study helps highlight the need to understand the cellular and genetic mechanisms that underpin a phenotype to fully understand its evolution and the power of evolution to manipulate mechanisms. Cytosine methylation regulates gene expression of many eukaryotes (Schmitz et al. 2019). Insects maintain cytosine methylation within gene bodies (Duncan et al. 2022). To begin it was parsimonious to assume that cytosine methylation also regulated gene expression of insects. However, careful molecular natural history has exposed much variation across insects for the level of gene body methylation and that differences of methylation are not casually associated with differences of gene expression. A change of mechanism addresses why cytosine methylation has persisted for insects and why it is not associated with gene expression across the taxon. Although the field does not have a fully supported explanation for why some but not all insects both retain cytosine methylation and its direct mechanism of action, it is clear that the explanation of cytosine methylation cannot be borrowed from many other branches of the tree-of-life to explain the function of insect cytosine methylation.

## Supporting information

RNA-seq library metadata

GO Annotations

7-day DEGs

14-days DEGs

GO terms 7-days DEGs

GO terms 14-days DEGs

GO terms Bewick et al 2019 Overlap

Supplementary Files Summary

## Acknowledgements

We appreciate the discussions and contributions from other members of our laboratory, especially Ahva Potticary. This work was funded by the USDA-ARS Non-Assistance Cooperative Agreement “Managing whiteflies and whitefly-transmitted viruses in vegetable crops in the southeastern U.S.” (#58-6080-9-006) to AJM and support from the National Science Foundation (MCB-1856143) to RJS.

## Author Contribution Statement

CBC, RJS, AJM, and PJM designed the study. CBC, EJS, ECM, and PJM preformed the experiments and analyzed the data. CBC, RJS, PJM and PJM provided interpretation of the data. CBC drafted the initial manuscript and authors contributed to subsequent revisions.

## Data Archiving

All high-throughput data is available under NCBI BioProject # XXXX. Phenotypic data for spermatocytes can be found on Dryad # XXXX.

## References

Alexa A, Rahnenfuhrer J, Lengauer T (2006) Improved scoring of functional groups from gene expression data by decorrelating GO graph structure. Bioinformatics 22, 1600–1607.

Arsala D, Wu X, Yi SV, Lynch JA (2022) *Dnmt1a* is essential for gene body methylation and the regulation of the zygotic genome in a wasp. PLoS Genetics 18:e1010181.

Amukamara AU, Washington JT, Sanchez Z, McKinney EC, Moore AJ, Schmitz RJ et al. (2020) More than DNA methylation: does pleiotropy drive the complex pattern of evolution of *Dnmt1*? Frontiers Ecol Evol 8, 10.3389.

Andrews S (2010) FastQC: A quality control tool for high throughput sequence data. http://www.bioinformatics.babraham.ac.uk/projects/fastqc/

Benjamini Y, Hockberg Y (1995) Controlling the false discovery rate: A practical and powerful approach to multiple testing. J R Stat Soc 57:289–300.

Bewick AJ, Vogel KJ, Moore AJ, Schimitz RJ (2017) Evolutin of DNA methylation across insects. Molec Biol Evol 34:654–665.

Bewick AJ, Sanchez Z, McKinney EC, Moore AJ, Moore PJ, Schmitz RJ (2019) *Dnmt1* is essential for egg production and embryo viability in the large milkweed bug *Oncopeltus fasciatus*. Epigenetics & Chromatin 12:1–14.

Cantalapiedra CP, Hernández-Plaza A, Letunic I, Bork P, Huerta-Cepas J (2021) eggNOG-mapper v2: functional annotation, orthology assignments, and domain prediction at the metagenomic scale. Molec Biol Evol 38:5825–5829.

Chesebro J, Hrycaj S, Mahfooz N, Popadic’ A (2009) Diverging functions of *scr* between embryonic and postembryonic development in a hemimetabolous insect, *Oncopeltus fasciatus*. Develop Biol 329:142–151

Chipman AD (2017) *Oncopeltus fasciatus* as an evo-devo research organism. Genesis 55:e23020.

Cunningham CB, Douthit MK, Moore AJ (2014) Octopaminergic gene expression and flexible social behaviour in the subsocial burying beetle *Nicrophorus vespilloides*. Insect Molec Biol 23:391–404.

Cunningham CB, Ji L, McKinney EC, Benowitz KM, Schmitz RJ, Moore AJ (2019) Changes of gene expression but not cytosine methylation are associated with male parental care reflecting behavioural state, social context and individual flexibility. J Exp Biol 222:jeb188649.

Duncan EJ, Cunningham CB, Dearden PK (2022) Phenotypic plasticity: what has DNA methylation got to do with it? Insects 13:110.

Gegner J, Gegner T, Vogel H, Vilcinskas A (2020) Silencing of the *DNA methyltransferase 1 associated protein 1 (DMAP1)* gene in the invasive ladybird *Harmonia axyridis* implies a role of the *DNA methyltransferase 1-DMAP1* complex in female fecundity. Insect Molec Biol 29:148–159.

Glastad KM, Goodisman MA, Yi SV, Hunt BG (2016) Effects of DNA methylation and chromatin state on rates of molecular evolution in insects. G3: Genes Genomes Genetics 6, 357–363.

Hans F, Dimitrov S (2001) Histone H3 phosphorylation and cell division. Oncogene 20:3021–3027.

Huang DW, Sherman BT, Lempicki RA (2009) Bioinformatics enrichment tools: paths toward the comprehensive functional analysis of large gene lists. Nucleic Acids Res 37:1–13.

Kassambara A, Mundt F (2020) Factoextra: Extract and visualize the results of multivariate data analyses. v1.0.7. CRAN.R-project.org/package=factoextra

Kim D, Paggi JM, Park C, Bennett C, Salzberg SL (2019) Graph-based genome alignment and genotyping with HISAT2 and HISAT-genotype. Nature Biotech 37:907–915.

Kopylova E, Noé, Touzet H (2012) SortMeRNA: Fast and accurate filtering of ribosomal RNAs in metatranscriptomic data. Bioinformatics 28:3211–3217.

Korotkevich F, Sukhov V, Budin N, Shpak B, Artyomov MN, Sergushichev A (2021) Fast gene set enrichment analysis. bioRxiv, doi.org/10.1101/060012.

Law JA, Jacobsen, SE (2010) Establishing, maintaining, and modifying DNA methylation patterns in plants and animals. Nature Rev Genetics 11:204–220.

Libbrecht R, Oxley PR, Keller L, Kronauer DJC (2016) Robust DNA methylation in the clonal raider ant brain. Curr Biol 26:391–395.

Livak KJ, Schmittgen TD (2001) Analysis of relative gene expression data using real-time quantitative PCR and the 2(-Delta Delta C(T)) method. Methods 25:402–408.

Love MI, Huber W, Anders S (2014) Moderated estimation of fold change and dispersion for RNA-seq data with DESeq2. Genome Biolo 15:550.

Magoc T, Salzberg S (2011) FLASH: fast length adjustment of short reads to improve genome assemblies. Bioinformatics 27:2957–2963

Martin M (2011) Cutadapt removes adapter sequences from high-throughput sequencing reads. EMBnet Journal, 17, 10–12.

Panfilio KA, Jentzsch IMV, Benoit JB, Erezyilmaz D, Suzuki Y, Colella S, et al. (2019) Molecular evolutionary trends and feeding ecology diversification in the Hemiptera, anchored by the milkweed bug genome. Genome Biol 20:64.

Pertea M, Pertea GM, Antonescu CM, Chang T-C, Mendell JT, Salzberg S (2015) StringTie enables improved reconstruction of a transcriptome from RNA-seq reads. Nature Biotechnol 33:290–295.

Prigent C, Dimitrov S (2003) Phosphorylation of serine 10 in histone H3, what for? J Cell Sci 116:3677–3685.

Provataris P, Meusemann K, Neihuis O, Garth S, Misof B (2018) Signatures of DNA Methylation across insects suggest reduced DNA methylation levels in holometabola. Genome Biol Evol 10:1185–1197.

R Core Team (2021) R: A language and environment for statistical computing. R Foundation for Statistical Computing, Vienna, Austria. https://www.R-project.org/.

RStudio Team (2022) RStudio: Integrated Development for R. RStudio, PBC, Boston, MA URL http://www.rstudio.com/.

Schmitz RJ, Lewis ZA, Goll MG (2019) DNA methylation: Shared divergent features across eukaryotes. Trends in Genetics 35:818–827.

Schulz NKE, Wagner CI, Ebeling J, Raddatz G, Diddens-de Buhr M, Lyko F et al. (2018) *Dnmt1* has an essential function despite the absence of CpG DNA methylation in the red flour beetle *Tribolium castaneum*. Sci Reports 8, 16462.

Soneson C, Love MI, Robinson MD (2015) Differential analyses for RNA-seq: transcript-level estimates improve gene-level inferences. F1000Research, f1000research.7563.1.

Stephens M (2016) False discovery rates: a new deal. Biostatistics, 18:2.

Supek F, Bošnjak M, Škunca N, Šmuc T (2011) REVIGO summarizes and visualizes long lists of Gene Ontology terms. PLoS ONE 6, e21800.

Magoc T, Salzberg S (2011) FLASH: Fast length adjustment of short reads to improve genome assemblies. Bioinformatics 27:2957–63.

Maleszka R, Kucharski R (2022) Without mechanism, theories and models in insect epigenetics remain a black box. Trends in Genetics 38:1108–1111.

Martin M (2011) Cutadapt removes adapter sequences from high-throughput sequencing reads. EMBnet.journal 17:10–12.

Morandin C, Brendel VP (2021) Tools and applications for integrative analysis of DNA methylation in social insects. Mole Ecol Res 22:1656–1674

Panfilio KA, Vargas Jentzsch IM, Benoit JB et al. (2019) Molecular evolutionary trends and feeding ecology diversification in the Hemiptera, anchored by the milkweed bug genome. Genome Biol 20:64.

Pertea M, Pertea GM, Antonescu CM, Chang TC, Mendell JT, Salzberg SL (2015) StringTie enables improved reconstruction of a transcriptome from RNA-seq reads. Nature Biotech 33:290–295.

Provataris P, Meusemann K, Niehuis O, Grath S, Misof B (2018) Signatures of DNA methylation across insects suggest reduced DNA methylation levels in holometabola. Genome Biol Evol 10:1185–1197.

Washington JT, Cavender KR, Amukamara AU, McKinney EC, Schmitz RJ, Moore PJ (2021) The essential role of *Dnmt1* in gametogenesis in the large milkweed bug Oncopeltus fasciatus. eLife 10:e62202.

Yan H, Bonasio R, Simola DF, Liebig J, Berger SL, Reinberg D (2014) DNA methylation in social insects: how epigenetics can control behavior and longevity. Annu Rev Entomol 60:435–452.

Zhu A, Ibrahim JG, Love MI (2018) Heavy-tailed prior distributions for sequence count data: removing the noise and preserving large differences. Bioinformatics 35:2084–2092.

Zwier MV, Verhulst EC, Zwahlen RD, Beukeboom LW, van de Zande L (2012) DNA methylation plays a crucial role during early *Nasonia* development. Insect Molec Biol 21:129–138.

