## Supplementary Files Summary for "The critical role of *Dmnt1* during spermatogenesis is not predictable, but knockdown does cause pervasive differential transcription"

Supplementary File Summaries

Supplementary File 1 - SI_FIle_1_Library_Metadata_v1.txt. Contains the for the RNA-seq libraries used here.

Supplementary File 2 - SI_FIle_2_GO_annotations_v1.txt. Contains the updated GO term annotations for *Oncopeltus fasciatus*.

Supplementary File 3 - SI_File_3_Onco_testes_Day7_DEGs_v1. Contains the output from DESeq2 for the differential gene expression of the 7-day samples.

Supplementary File 4 - SI_File_4_Onco_testes_Day14_DEGs_v1. Contains the output from DESeq2 for the differential gene expression of the 14-day samples.

Supplementary File 5 - SI_FIle_5_GO_Overrepresentation_Day7_v1. Contains the output from topGO for the GO term enrichment analysis of the 7-day samples.

Supplementary File 6 - SI_FIle_6_GO_Overrepresentation_Day14_v1. Contains the output from topGO for the GO term enrichment analysis of the 14-day samples.

Supplementary File 7 - SI_FIle_7_GO_Overrepresentation_Bewick-Overlap-14_v1. Contains the output from topGO for the GO term enrichment analysis of the 14-day samples.
